# Glomerulus-Targeted Nanotherapy via Collagen IV-α3 Binding Enhances Renal Immunoregulation in Lupus Nephritis

**DOI:** 10.64898/2026.07.14.738533

**Authors:** Lei Wang, Eleni Markoutsa, Rajkumar Venkatadri, Rahul Sharma, Carlos Esquivel, Akshata Patne, Jin Wei, Jie Zhang, Kristof Williams, William Lawless, Amtul Muskan, Liying Fu, Karthick Mayilsamy, Subhra Mohapatra, Shyam Mohapatra, Ruisheng Liu

**Affiliations:** Department of Molecular Pharmacology and Physiology, University of South Florida College of Medicine, Tampa, FL 33612, USA; Laboratory for Translational Nanotechnology, Department of Internal Medicine, University of South Florida, Tampa, FL, 33612, USA; Center for Immunity Inflammation and Regenerative Medicine, Division of Nephrology, University of Virginia School of Medicine, Charlottesville, VA 22903, USA; Division of Nephrology, Department of Medicine, Chobanian & Avedisian School of Medicine, Boston University, Boston, MA 02118, USA; Department of Molecular Medicine, University of South Florida, Tampa, FL, 33612, USA; James A Haley VA Hospital, Tampa, FL, 33612, USA; Department of Laboratory Medicine and Pathology, Mayo Clinic Arizona, Scottsdale, AZ 85259, USA

**Keywords:** Glomerular basement membrane, Collagen IV (Col4-α3), Targeted drug delivery, Lupus nephritis, Nanoparticles

## Abstract

Lupus nephritis requires long-term immunosuppressive therapy, which is often associated with severe systemic side effects. Therefore, new therapeutic strategies that maintain high efficacy while minimizing adverse effects is essential. Although nanomedicine has advanced systemic and kidney-targeted drug delivery, a reliable method for glomerulus-specific delivery is lacking. Collagen IV (Col4)-alpha 3, located in the glomerular basement membrane (GBM) at the blood–tissue interface through fenestrated capillary endothelium, represents an ideal target for glomeruli delivery. Herein, we developed a novel liposomal nanoparticle conjugated with a Col4-alpha 3 binding peptide (Col4-α3-NPs) for selective glomerular targeting. Prednisolone-loaded Col4-α3-NPs were administered to lupus-prone mice twice weekly for 8 weeks. Kidney injury and function were evaluated biweekly, and renal immune cell populations were analyzed by flow cytometry at study completion. The results show that rhodamine-labeled NPs predominantly accumulate in kidney glomeruli 48 hours after intravenous injection. The Col4-NP system demonstrated stable and prolonged release of the encapsulated drug for over 48 hours. Lupus-prone mice treated with prednisolone-loaded Col4-NPs showed significantly improved renal function and histology, including a 30% increase in glomerular filtration rate (GFR), a 56% reduction in proteinuria, and decreased IgG deposition and fibrosis. Notably, treatment also enhanced renal regulatory T cell (Treg) populations. These findings suggest that glomerulus-targeted Col4-α3-NPs hold significant translational promise. This platform may offer an effective, site-specific treatment for lupus nephritis while minimizing systemic side effects and could be adapted for other glomerular diseases requiring targeted therapy.

## Introduction

Systemic lupus erythematosus (SLE) is a chronic autoimmune disorder characterized by the loss of immune tolerance and the production of autoantibodies, particularly antinuclear antibodies (1, 2). These autoantibodies form immune complexes that deposit in multiple tissues and organs, leading to widespread inflammation and organ dysfunction. Among the most serious complications of SLE is lupus nephritis (LN), which affects nearly 50% of patients and greatly increases the risk of end-stage renal disease (ESRD) as well as overall morbidity and mortality (3–5). LN primarily involves immune complex-mediated injury to the glomeruli, particularly the glomerular basement membrane (GBM) and mesangium (6–9). Current treatment of LN relies heavily on systemic immunosuppressive therapy—especially glucocorticoids (GCs)—to control inflammation and preserve kidney function (10, 11). However, these therapies are associated with substantial adverse effects, including increased susceptibility to infections, metabolic disturbances, and damage to non-target organs (12–14). This underscores a pressing need for targeted therapeutic strategies that minimize systemic exposure while delivering effective treatment directly to renal tissues.

Nanotechnology-based drug delivery systems offer a promising solution for organ-specific therapy. Liposomes, due to their biocompatibility and ability to encapsulate a wide range of therapeutic agents, have been extensively studied as drug carriers (15–17). However, traditional liposomes lack tissue specificity (18–20). To overcome this challenge, targeted nanoparticles have been developed for improved bioavailability and kidney glomerulus-specific accumulation (21–23).

Numerous preclinical studies have explored kidney-targeted nanoparticle systems (24–26), leveraging particle size, charge, and conjugation with targeting ligands to achieve selective accumulation in renal structures such as the GBM and mesangial cells (27–29). A variety of targeting peptides and antibodies have been utilized with nanoparticles trying to target the kidney and kidney cells (30–36). Despite some success, most of these approaches suffer from non-specific binding and lack durable, site-specific drug release.

We previously used the chitosan-Tripolyphosphate nanoparticles (chitosan-TPP NPs) encapsulated by liposomes exhibited excellent characteristics in drug delivery with controlled release (37). However, this complex is not a targeted system and does not selectively deliver drugs to the glomeruli in the kidneys. In this study, we developed a novel glomerulus-targeted drug delivery system by conjugating liposomal NPs with peptides specific to the α3 isoform of collagen IV (Col4-α3), which is highly expressed in the GBM — the only location in the entire body where Col4-α3 is directly exposed to circulating blood due to the fenestrated glomerular endothelium (38, 39). These Col4-α3-targeted and prednisolone-loaded liposomal NPs are designed to accumulate specifically in the glomeruli, where they stably and sustainably release their therapeutic payload. We loaded this system with prednisolone to assess its therapeutic efficacy in a lupus-prone mouse model.

Our results demonstrate that the Col4-α3–NP system selectively targets the glomeruli. Prednisolone-loaded Col4-α3–NP administered during the early phase of LN effectively attenuates renal injury and inflammation, preserves kidney function, and promotes the induction of local tolerogenic dendritic cells (DCs) and regulatory T cells (Tregs). The use of biocompatible and biodegradable components, combined with a clinically approved therapeutic agent, supports the translational potential of this platform. In addition to its therapeutic utility, this system holds promise as a diagnostic tool for in vivo imaging and assessment of nephron function.

## Materials and Methods

### Animals

Male C57BL/6J mice (8 weeks old) and female MRL-lpr mice (8 weeks old) were obtained from the Jackson Laboratory. After arrival, the animals were housed in a temperature-controlled environment with 12-hour light/12-hour dark cycle and ad libitum access to water and normal chow diet food (Teklad 2018C) for 1 week before experiments. Mice were randomly assigned into control or treatment groups. All procedures and experiments were performed at the animal facility of the University of South Florida Morsani College of Medicine with approval from the Institutional Animal Care and Use Committee of the University of South Florida (IACUC protocol number: IS00011754) and in accordance with all applicable federal guidelines. Animals were anesthetized with 2% isoflurane via inhalation for agent injections. At the end of the experiment, animals either underwent systemic saline perfusion through the left ventricle for organ or tissue collection under anesthesia or were euthanized by carbon dioxide asphyxiation followed by cervical dislocation, in accordance with the guidelines of the American Veterinary Medical Association. All chemicals were purchased from Sigma-Aldrich unless otherwise indicated.

### Surface Plasmon Resonance (SPR)

SPR was employed for binding measurements using Collagen Type I (US Biological) or Collagen Type IV Alpha 3 (COL4A3; RayBiotech) human recombinant protein. A GE Healthcare Biacore T200 was equipped with a CM5 chip. An average of approximately 5,000 RU of Collagens I and IV were crosslinked via Amine coupling chemistry via a contact time of 2000 sec and a flow rate of 5 μL/min. The proteins of interest were titrated and flowed at 30 μL/min in 1X phosphate buffered saline with surfactant P20 (PBS-P; pH 7.4) for 30 sec association time followed by a 120 sec dissociation. The sensorgrams were analyzed using Biacore T200 Software 3.0 and steady state was measured at 4 sec before injection stop, exported into Graphpad, and fit versus concentration using a one site specific binding model to calculate the apparent equilibrium dissociation constant (K_D_). Where appropriate, kinetics were measured using a 1:1 Langmuir binding model with Rmax set to local to obtain the association rate (K_on_), dissociation (K_off_), and the K_D_.

### Synthesis and characterization of liposomes

Liposomes consisted of 1,2-Distearoyl-sn-glycero-3-phosphocholine (DSPC), Cholesterol and 1,2-Distearoyl-sn-glycero-3-phosphoethanolamine-Polyethylene Glycol (DSPE-PEG)-maleimide at a molar ratio of 2:1:0.08 (Avanti Polar Lipids) were prepared using the thin film method as previously described (40). Briefly, the film was hydrated with pH 7.4 PBS-2mM EDTA and the liposomal size was reduced using probe sonication followed by extrusion through 100 nm pore size polycarbonate membranes. For the loading of prednisolone sodium phosphate into the Col4-α3-NPs the thin film was hydrated with PSP at a concentration of 92 mg/mL in water. Then, liposomes were incubated for 1 h at 55°C in order to anneal liposome structural defects. For the peptide conjugation onto liposomal surface, the peptide was first reduced using Tris(2-carboxyethyl)phosphine (TCEP) solution, at a 1:1 peptide:TCEP molar ratio. 0.21 mg of reduced peptide was added per mg of total lipid. The non-conjugated peptide was removed by dialysis. The peptide conjugation yield was estimated by Bradford and the lipid conjugation by Stewart assay. Size and Zeta potential were estimated by Dynamic Laser Scattering and TEM images were obtained by negative staining with 3% uranyl acetate. For the biodistribution and cell uptake studies the liposomal bilayer was labeled with 1,2-Distearoyl-sn-glycero-3-phosphoethanolamine (DSPE)-Rhodamine.

### Nanoparticle integrity

To investigate the integrity of the nanoparticles, calcein-loaded nanoparticles were prepared as described previously (40). In brief, the thin film was hydrated with 100 mM calcein (pH 7.4, adjusted to an osmolarity of 300 mosm). Non-encapsulated calcein was removed by size exclusion chromatography using a Sephadex column. Calcein was selected as a model drug due to its hydrophilic nature and molecular weight similar to that of prednisolone. Non-targeted and collagen-targeted nanoparticles, encapsulating calcein, were incubated at 37°C in PBS or 80% FBS, shaking at a final lipid concentration of 1 mg/mL. The percentage release of calcein at various time intervals was estimated by adding 10 µL of the samples to 1 mL of PBS or to 1 mL of PBS with 2% Triton-X (to determine the total fluorescence of the encapsulated calcein) and measuring the fluorescence at an excitation wavelength of 425 nm and an emission wavelength of 528 nm. The percentage release was calculated as the fluorescence intensity measured in PBS divided by the fluorescence intensity in PBS with Triton X, multiplied by 100.

### The uptake of the Col4-**α**3-NPs

To investigate the uptake mechanism of Col4-α3-NPs, primary cultured human umbilical vein endothelial cells (HUVECs; PCS-100-010, ATCC) were seeded in 96-well plates at a density of 5,000 cells per well and maintained in Vascular Cell Basal Medium (ATCC, PCS-100-030) supplemented with the Endothelial Cell Growth Kit—VEGF (ATCC, PCS-100-040). After 48 hours of culture, the cells were washed four times with fresh medium and incubated with Rhodamine-labeled, FITC-steroid–loaded Col4-α3-NPs (200 µg/ml) for 4 hours. Following incubation, cells were washed thoroughly with medium, and nanoparticle uptake was visualized using a laser scanning confocal microscope at 24-and 48-hours post-treatment.

### Transmission electron microscopy (TEM) to examine NP bindings

C57BL/6 mice received a single intravenous injection of gold nanoparticle-loaded Col4-α3-targeted liposomes at a dose of 0.3 mL/kg under 2% inhalational isoflurane anesthesia. Non-targeted nanoparticles (NPs) were used as controls. Twenty-four hours after injection, mice were re-anesthetized with isoflurane and systemically perfused with saline through the left ventricle. Then kidneys were removed and prepared for examination for NP bindings with TEM (JEM-1400Plus) as we described previously (41).

### Drug release characteristics of the Col4-**α**3-NP system

To investigate the release of prednisolone from nanoparticles, we developed a high-performance liquid chromatography (HPLC) method. The HPLC analysis was conducted using a Shimadzu system with separation achieved at 40 °C using a Thermo Scientific™ Hypersil GOLD™ PFP column. The analyte was eluted with a mobile phase consisting of acetonitrile and water in a 25:75 (v/v) ratio, at a flow rate of 1 mL/min. Prednisolone was detected at 245 nm with a retention time of approximately 15 min.

For release studies, prednisolone-loaded Col4-α3-NPs were prepared as previously described (42). Non-encapsulated prednisolone was removed through dialysis. To study the prednisolone release in vivo, C57BL/6 mice were injected with either free prednisolone (6 mg/kg) or an equivalent dose encapsulated in Col4-α3-NPs under 2% inhalational isoflurane anesthesia. Plasma samples were collected at 4, 10, and 24h post-injection. Prednisolone levels were quantified by HPLC as described above and previously reported (43).

### Biodistribution of the Col4-**α**3-NPs

C57BL/6 male mice were injected intravenously (i.v.) with rhodamine-or DiR-labeled NPs at a dosage of 40 mg/kg of lipid under 2% inhalational isoflurane anesthesia, either with or without Col4-α3 targeting. Forty-eight hours after injection, the mice were re-anesthetized with 2% isoflurane and systemically perfused with saline through the left ventricle. The main high accumulation organs including liver, spleen and kidney were obtained for fluorescence imaging with IVIS. The radiant efficiency, defined as fluorescence intensity/area/time, was analyzed using the IVIS imaging software with the region of interest (ROI) tool. To highlight kidney-selective accumulation of Col4-α3-NPs, fluorescence signal intensity was quantified after background subtraction and expressed as the ratio of Col4-α3-NPs to NT-NPs for each organ.

To detect the specific localization of the NPs in the kidneys, C57BL/6 male mice were injected intravenously (i.v.) with DiR-labeled NPs at a dosage of 40 mg/kg of lipid, either with or without Col4-α3 targeting. Forty-eight hours after injection, the perfused kidneys were slightly fixed in 4% PFA, sectioned to a thickness of 100 μm using a vibratome (Leica Model 1000), and stained for nuclei with DAPI. The samples were visualized using a Keyence BZ-X 700 fluorescence microscope.

### Assessment of renal function and hemodynamic impact of nanoparticles

To assess whether the administered nanoparticles, with or without Col4 targeting, impact renal function, hemodynamics, or other physiological parameters, we performed functional evaluations in conscious male C57BL/6J mice following intravenous injection of either Col4-α3-NPs or NT-NPs at a dose of 40 mg/kg under 2% inhalational isoflurane anesthesia. At 24 hours post-injection, blood samples were collected for blood gas analysis, including pH, pCO₂, pO₂, bicarbonate, and lactate levels. Renal function was assessed by measuring GFR using a transcutaneous FITC-sinistrin clearance method as previously described (44, 45). Additionally, kidney injury markers such as plasma creatinine (PCr) and blood urea nitrogen (BUN) were quantified as previously described (45–47). Anesthetized blood pressure and renal blood flow were also measured using a Transonic flow probe (Model #: 0.5 PS) with the PowerLab software as previously described (48).

### Examination of the preventive therapeutic efficacy of prednisolone-loaded Col4-α3-NPs for lupus nephritis

SLE-prone female MRL-lpr mice (8 weeks old) were randomly assigned into experimental and control groups. Female MRL/lpr mice were used throughout the therapeutic efficacy studies because they develop spontaneous lupus nephritis with a higher disease incidence and more consistent progression than males, making them the appropriate model for evaluating treatment efficacy. Mice were injected i.v. twice a week for 8 weeks with either prednisolone-loaded Col4-α3-NPs (0.34 mg/kg) or NT-NPs under 2% inhalational isoflurane anesthesia. Plasma and urine were collected biweekly to measure Pcr and proteinuria. As we previously described (45, 49), urine samples were collected in metabolic cages for 24hours. Pcr was measured by LC-MS at the O’Brien Center, University of Alabama at Birmingham. Proteinuria concentration was measured with ELISA kit (Abcam) and daily albuminuria was calculated. GFR were measured by transcutaneous GFR monitoring (MediBeacon, MB 0309) in conscious mice every two weeks during the treatment as previously described (45–47). At the end of the 8-week treatment period, kidneys were harvested for histological and flow cytometry analyses.

### Histology

The other kidney was collected for histopathological analysis by PAS, Masson’s Trichrome staining and immunofluorescent staining with anti-IgG (ab150113; Abcam) to measure IgG deposition as we previously described (50–52). Five-Ten randomly chosen fields were captured, and the degree of injury and fibrosis were determined and quantified in each image as reported (41). The IgG deposition was visualized with a confocal microscope and quantified by the mean fluorescence of 50 glomerular cross sections from the respective mice using the Image J software (51–53).

### Flow cytometry analysis of immune cells

Flow cytometry on lymphoid and kidneys were performed utilizing the fluorophore-labeled antibodies as described before(47). Briefly, MRL-lpr mice were euthanized, and kidneys and renal lymph nodes were collected. Kidneys were chopped and digested in RPMI containing collagenase D (1 mg/mL) and DNase I (0.1 mg/mL) at 37°C for 30 minutes. Tissues were then passed through a 70-μm cell strainer, and red blood cells were lysed using ACK buffer. Lymph nodes were gently mashed through a cell strainer without enzymatic digestion. Cells were washed and resuspended in FACS buffer (PBS with 2% FBS) and incubated with anti-CD16/32 (Fc block) for 10 minutes at 4°C. Surface staining was performed using fluorochrome-labeled antibodies against CD45, CD11c, CD11b, MHC II (IA/IE), CD3, CD4, CD8, B220, CD69, F4/80, CD80, and CD86. For intracellular cytokine staining (e.g., IFN-γ, IL-17A, TNF-α), cells were stimulated with PMA and ionomycin in the presence of brefeldin A for 4 hours, followed by fixation and permeabilization. For Foxp3 staining, Foxp3 staining kit (eBiosciences) was used. Samples were run on a BD FACS Calibur flow cytometer and data were analyzed using FlowJo software (BD Biosciences). Immune cell subsets were identified based on gating of live, singlet CD45⁺ cells and lineage markers.

## Statistics

All data are presented as mean ± standard deviation (SD), except where otherwise noted. Differences between 2 groups were assessed by unpaired 2-tailed Student’s t test. The number of animals required to achieve a value of 0.05 and a 1-β value of 0.8 was predetermined on the basis of preliminary experiments. One-way or two-way ANOVA was used for comparisons among more than 2 groups and was followed by Bonferroni’s post hoc test. Statistical significance was set at p < 0.05. All analyses were performed using GraphPad Prism (version 9.0 h).

## Results

### Identification of Col4-α3 targeted peptide and evaluation of the Col4-α3-specific targeting

To identify Col4 targeted peptides, we utilized the crystal structure of Col4-α3 with Molecular Operating Environment software (MOE, Chemical Computing Group, Montreal, CA) and searched a membrane binding motif database, which led to the identification of several unique peptides that could bind specifically to Col4-α3. Figure 1A and B illustrate the ligand interaction residues involved in the binding interactions between the computationally designed peptide sequence (KLWVLPKGGGC) and the experimentally determined crystal structure of collagen type IV α3 NC1 domain from the native α3α4α5 NC1 hexamer (PDB: 5NB0), reflecting the trimer–trimer interface in the GBM collagen IV scaffold, as analyzed using Schrödinger’s Maestro software with the GLIDE and Ligand Interaction Diagram extensions. This structural model, paired with Surface Plasmon Resonance (SPR) validation on trimeric NC1 domains, ensures computational predictions align with physiological relevance.

**Figure 1.**
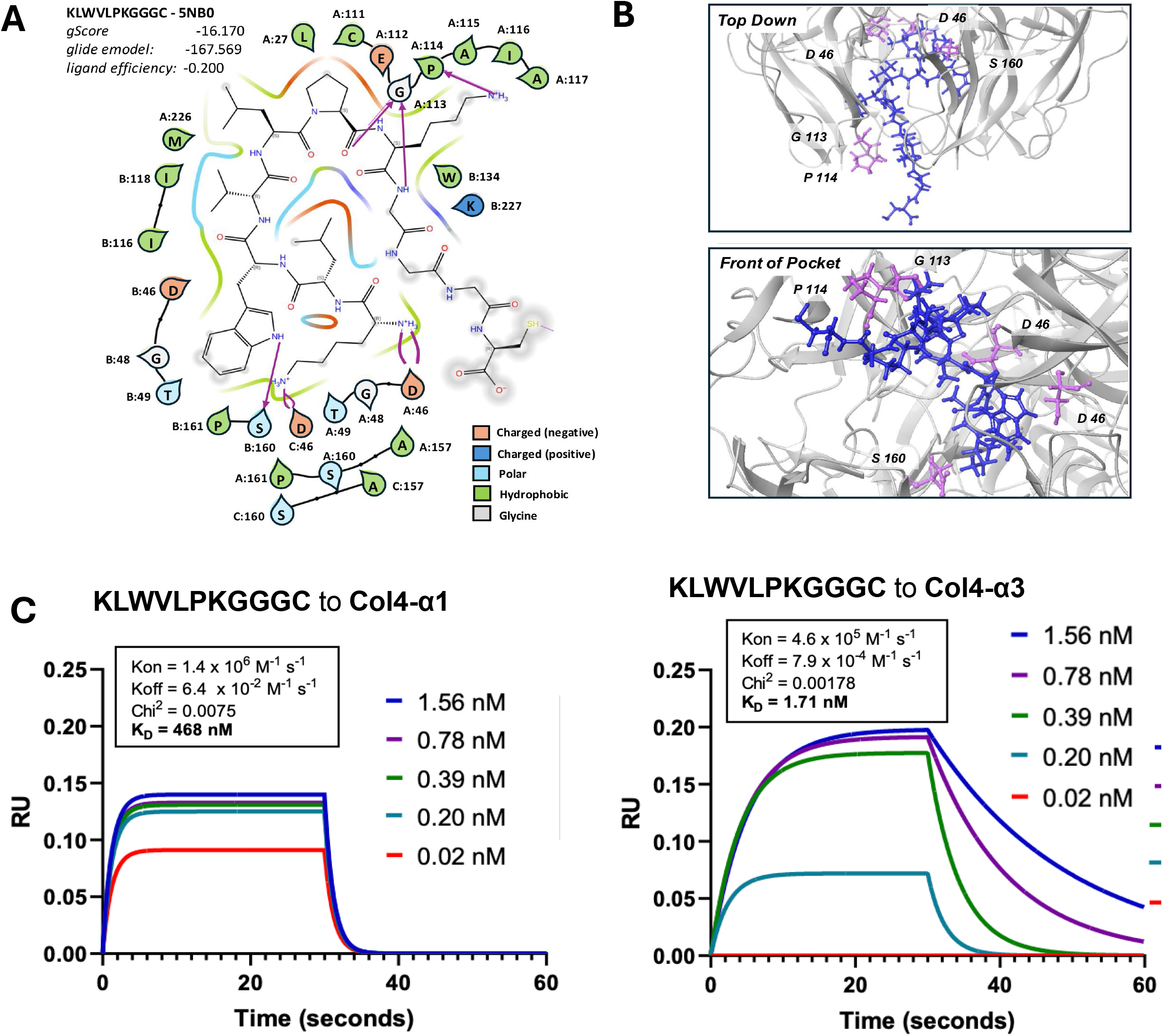
**Evaluation of the specific targeting of the COLIV-targeted peptide to Col4-**α**3. A**-**B**. 2D ligand interaction diagram identifying residues involved in binding interactions of the computational design model peptide sequence (KLWVLPKGGGC) on the experimentally determined crystal structure for collagen type IV alpha3NC1 (PDB: 5NB0). **C**. Surface Plasmon Resonance (SPR) to evaluate the binding affinity of the targeted peptide to Col4-α3 and the binding of Col4-α1 as negative control.

To evaluate the Col4-α3-specific binding of this peptide, we developed a SPR protocol for immobilizing Col4-α1 (negative control) and Col4-α3 on a CM5 sensor chip via a standard amine-coupling reaction. Affinity constants, including association rate (Kon, ka), dissociation rate (Koff, kd), and equilibrium dissociation constant (KD), were directly measured during the analysis of three custom-designed peptide fragments as previously described (54). In Figure 1C, the binding affinity of the targeted peptide to control Collagen Type I Alpha 1 (Col-α1) is shown. The peptide exhibited low binding affinity, with a K_on_ of 1.4 × 10□M⁻¹·s⁻¹, a K_off_ of 6.4 × 10⁻² M⁻¹·s⁻¹, and a K_D_ of 468 nM (Chi² = 0.0075). For Col1α1, the equilibrium K_D_ was determined by steady-state fitting due to non-ideal binding kinetics. The apparent K_off_/K_on_ ratio is not used as the primary K_D_ because of likely avidity and mass transport effects common in peptide–fibrillar collagen interactions. These results indicate a weak interaction between the identified peptide and Col-α1. In contrast, the peptide demonstrated a high binding affinity with Col4-α3, with a K_on_ of 4.6 × 10□M⁻¹·s⁻¹, a K_off_ of 7.9 × 10⁻□s⁻¹, and a K_D_ of 1.71 nM (Chi² = 0.00178). These studies validated high-affinity binding of select peptide sequence to Col4-α3.

### Preparation and characterization of Col4-**α**3-NPs loaded with prednisolone

Liposomes were prepared with DSPC, cholesterol, and DSPE-PEG-maleimide in a molar ratio of 2:1:0.08, as previously described (55, 56) (Figure 2A). The collagen-targeted peptide was conjugated to the liposomal surface via a C-terminal GGGC linker to DSPE-PEG-maleimide after being reduced with a TCEP solution, using a 1:1 molar ratio of peptide to TCEP (Figure 2B). The yield of peptide conjugation was estimated using a Bradford assay, while the lipid concentration was determined with the Stewart assay. The peptide conjugation yield was found to be 75%, which translates to approximately 1,000 peptide molecules per liposome. Transmission electron microscopy (TEM) images confirmed that the pegylated CoI4-targeted liposomes had a size of 100 nm and exhibited a spherical morphology (Figure 2C). Dynamic laser scattering was used to assess both size and Zeta potential, revealing that the pegylated Col4-NPs were slightly larger than the non-targeted particles (NT-NPs) (Figure 2D). High-performance liquid chromatography (HPLC) and the Stewart assay were utilized to estimate the ratio of prednisolone to lipid, as described in the methods section. The prednisolone to lipid ratio was found to be 1:5 weight/weight.

**Figure 2.**
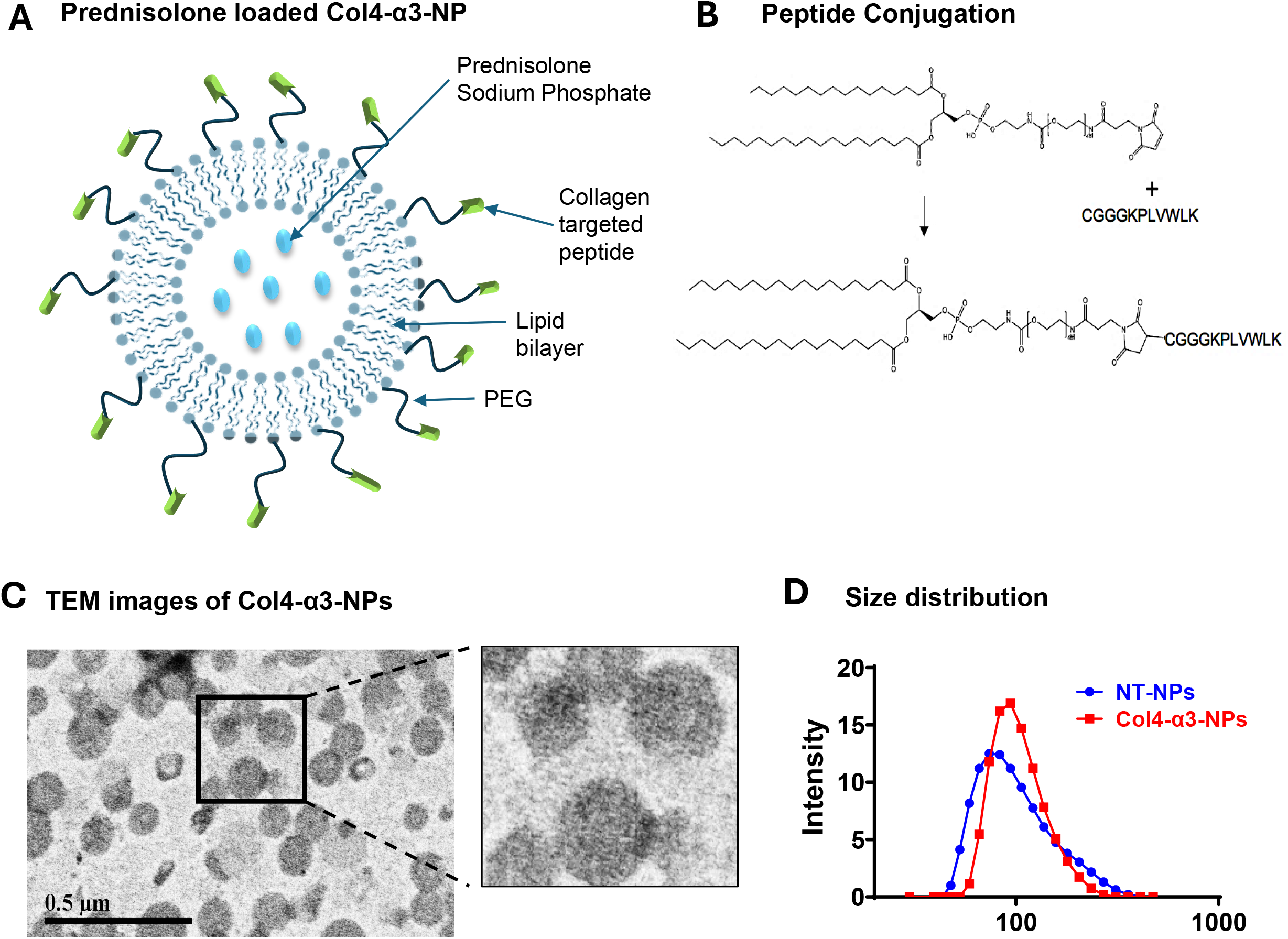
**Preparation and characterization of Col4-**α**3-NPs loaded with prednisolone. A**. Schematic representation of prednisolone loaded Col4-α3-NP system. **B**. Peptide conjugation. **C**. TEM images of Col4-α3-NPs **D**. Size distribution of non-targeted (NT-NPs) and Col4-α3-NPs.

### Uptake of Col4-**α**3-NPs

After synthesizing and physically characterizing the Col4-α3-NPs, we investigated their uptake mechanism. To study renal uptake, gold nanoparticles (5 nm in diameter) were incorporated into the Col4-α3-NPs. C57BL/6 male mice were intravenously (i.v.) injected with the gold-loaded NPs at a lipid concentration of 40 mg/kg. Kidney tissue was collected 24 hours post injection, fixed and sections observed under JEOL JEM-1400 TEM. As demonstrated in Figure 3A, the gold-loaded Col4-α3-NPs were detected within the endothelium of the glomerular capillaries.

**Figure 3.**
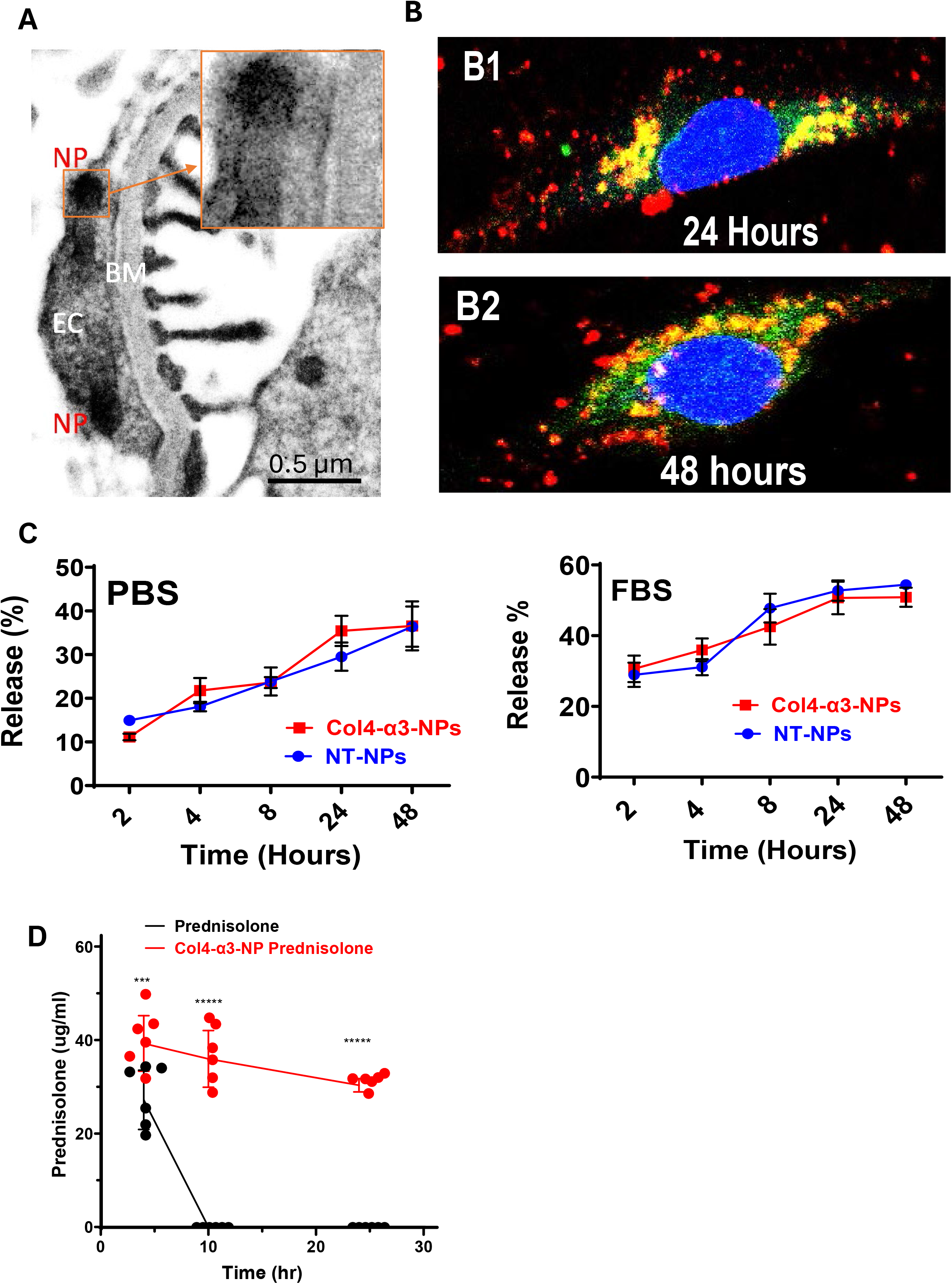
Targeted delivery and extended release of Liposomal prednisolone. **A**. TEM Imaging of kidney tissue of mice injected with Col4-α3-Gold NPs. Gold NPs (approximately 5 nm in diameter) loaded into Col4-α3-NPs were detected within the endothelium 24 hours after intravenous administration. (NP: nanoparticles; BM: Basement Membrane; EC: Endothelial Cell.) **B**. In vitro uptake of Col4-α3-NPs. Rhodamine-labeled and FITC-steroid-loaded Col4-α3-NPs were incubated with HUVECs for 4 hours. The signals were observed using a laser scanning confocal microscope at 24 and 48 hours after Col4-α3-NP treatment. **C**. Release of model drug (Calcein) from NT-NPS and Col4-α3-NPs in PBS and FBS. **D.** Plasma concentrations of prednisolone in mice receiving intravenous administration of 6 mg/kg of either free prednisolone or Col4-α3-NPs –encapsulated prednisolone were compared. Statistical analysis was performed using Two-way ANOVA followed by Tukey’s multiple comparisons test. ***P<0.001, ****p<0.0001, n=6 mice/group.

To demonstrate the retention of Col4-α3-NPs inside the cells, we used a dual labeling technique. The outer layer of the Col4-α3-NPs was labeled with Rhodamine-PE lipid (red), while the Col4-α3-NPs were loaded with a water-soluble FITC-labelled steroid (green). Human Umbilical Vein Endothelial Cells (HUVEC) at 70% confluency were treated with 2 μM of the dual-labeled Col4-α3-NPs for 4 hours, followed by three washes. Fluorescent images were captured at 24 and 48 hours using a laser scanning confocal microscope (Olympus FV1200). As shown in Figure 3B, the red and green signals were nearly completely colocalized at 24 hours, indicating that only a small percentage of the FITC-steroid had been released from the Col4-α3-NPs at that time point. By 48 hours post-treatment, more FITC-steroid had been released, confirming the slow intracellular drug release properties of the Col4-α3-NPs.

### Prolonged release of liposomal prednisolone

To investigate the release profile of hydrophilic drugs from both targeted and non-targeted liposomes, calcein was employed as a model drug. Calcein has a molecular weight and water solubility that are very similar to those of prednisolone sodium phosphate. A 100 mM calcein solution was encapsulated into the liposomes, and any excess was removed using a Sephadex G-50 column through size exclusion chromatography. When calcein is encapsulated in liposomes at high concentrations, it exhibits self-quenching, resulting in low fluorescence. The release of calcein from the liposomes causes a significant increase in fluorescence. As shown in Figure 3C, calcein release from both targeted and non-targeted liposomes in PBS was found to be 40% after 48 hours, indicating that the presence of the peptide on the surface of the liposomes does not affect their integrity. Furthermore, the release of calcein from both types of liposomes in the presence of serum proteins did not show any significant differences, suggesting that there is no interaction between the peptide and the serum proteins that could compromise the integrity of the liposomes.

To evaluate the prolonged release of prednisolone from NPs, we measured plasma prednisolone concentrations in mice following intravenous administration of 6 mg/kg of either free prednisolone or Col4-α3-NPs–encapsulated prednisolone. Figure 3D displays the mean plasma prednisolone concentrations at 4 hours, 10 hours, and 24 hours post injection for both formulations. Free prednisolone was completely eliminated from the plasma by 10 hours post-administration. In contrast, liposomal prednisolone remained in the plasma much longer, with concentrations exceeding 30 µg/ml still detectable at 24 hours post-injection.

### Biodistribution of Col4-**α**3-NPs

To evaluate nanoparticle biodistribution, C57BL/6 mice were intravenously injected with rhodamine-labeled NT-NPs or Col4-α3-NPs. Organs were harvested 48 hours post-injection, and fluorescence signals were visualized using an in vivo imaging system (IVIS). As shown in Figures S1A and S1B, the Col4-α3-NP group exhibited the highest signal intensity in the kidneys at 48 hours post-injection, whereas fluorescence signals in the liver and spleen were low and comparable between the targeted and non-targeted groups. Signals in the heart and lungs were negligible in all conditions. Because photobleaching of rhodamine may affect signal intensity over time, nanoparticle biodistribution was further confirmed using DiR-labeled NT-NPs or Col4-α3-NPs. C57BL/6 mice were intravenously injected, organs were harvested 48 hours post-injection, and fluorescence signals were analyzed by IVIS. As shown in Figures 4A and 4B, the Col4-α3-NP group displayed the highest signal intensity in the kidneys, with approximately twofold greater fluorescence compared with NT-NP–treated kidneys, confirming enhanced renal accumulation of Col4-α3-NPs.

**Figure 4.**
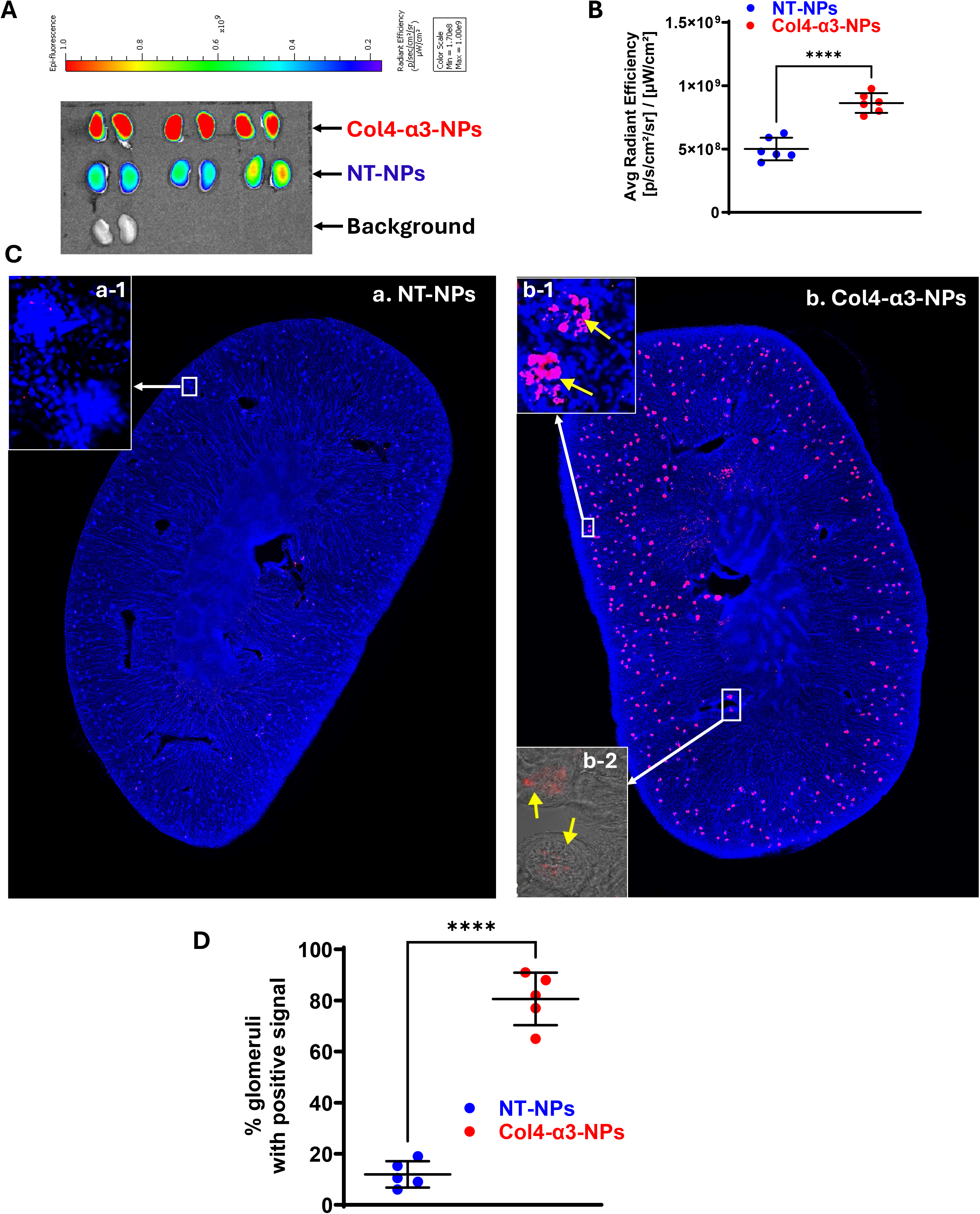
**Biodistribution of Col4-**α**3-NPs. A** Representative image of mouse kidneys collected 48 hours after injection with DIR-labeled NT-NPs or Col4-α3-NPs. **B**. Quantification of fluorescence signal intensity, expressed as the ratio of Col4-α3-NPs to NT-NPs kidneys. **C**. Fluorescent images of kidney sections after 48 hours of intravenous injection of DiR-labelled NT-NPs (a) and Col4-α3-NPs (b). Images a1, b1 and b2 show higher-magnification views of glomeruli. In image b2, the overlay of a bright-field image with the DiR fluorescein signal at high magnification confirms localization of the red signal to the glomerular area. D. Quantitative analysis of the percentage of glomeruli exhibiting positive fluorescence signals. Statistical analysis was performed using unpaired 2-tailed Student’s *t* test. ****P<0.0001, n=4 mice/group.

To further confirm glomerular-specific binding of the targeted nanoparticles, C57BL/6 mice were injected with DiR-labelled NT-NPs or Col4-α3-NPs. Kidneys were collected at 48 hours, lightly fixed, and sectioned for fluorescence microscopy. As shown in Figure 4C, DiR-labeled Col4-α3-NPs (red) were prominently localized within the renal cortex, while NT-NPs showed minimal detectable signal. Under identical imaging conditions, DiR-labeled Col4-α3-NPs were clearly and specifically visualized within the renal glomeruli (panel b), whereas NT-NPs showed limited detectable glomerular localization in any region of the kidney (panel a). Figure 4D summarizes data from five mice, showing that over 80% of glomeruli in the Col4-α3-NPs kidneys exhibited a positive Dir signal, whereas less than 10% of glomeruli in the NT-NPs kidneys showed detectable signals. Together, these findings demonstrate that Col4-α3-NPs preferentially accumulate in kidney glomeruli with minimal off-target distribution, indicating effective and selective glomerular targeting.

### Col4-NPs have no adverse effects on kidney function or hemodynamic parameters

To evaluate whether NPs, with or without Col4-α3 targeting, affect renal hemodynamics, function, or physiological parameters, C57BL/6 mice were intravenously injected with the respective NPs. Blood, urine, and tissue samples were collected and analyzed as described in the Methods. As shown in Figure S2 and Table S1, neither targeted nor non-targeted NPs had significant effects on kidney function. Plasma creatinine, Blood Urea Nitrogen (BUN), Glomerular Filtration Rate (GFR), and Renal Blood Flow (RBF) remained unchanged following NP administration. Additionally, Periodic Acid–Schiff (PAS) staining revealed no morphological alterations in kidney tissue (Fig. S2.D), confirming the absence of NP-induced renal injury.

### Targeted delivery of prednisolone via Col4-**α**3-NPs preserves renal function in MRL-lpr Mice

The primary objective of this study was to determine whether glomerular-targeted nanoparticle delivery could achieve therapeutic efficacy with markedly reduced glucocorticoid exposure. Since the amount of prednisolone encapsulated in the nanoparticles (0.34mg/kg) was approximately 20-fold lower than the conventional dose (5-10mg/kg)(57–60) used for lupus treatment, we first performed a preliminary study in female MRL-lpr mice using the same low dose of free prednisolone and empty NPs as controls (Figure S3). Neither treatment produced meaningful therapeutic benefit. We therefore evaluated the efficacy of prednisolone-loaded Col4-α3-NPs at this low dose. Female MRL-lpr mice received intravenous injections of Col4-α3-NPs or NT-NPs loaded with a low dose of prednisolone (0.34 mg/kg) twice weekly for eight weeks. Kidney function and injury were assessed biweekly, as illustrated in Figure 5A. At baseline, plasma creatinine, proteinuria and GFR were comparable between the two groups, with no significant tissue injury observed at treatment initiation.

**Figure 5.**
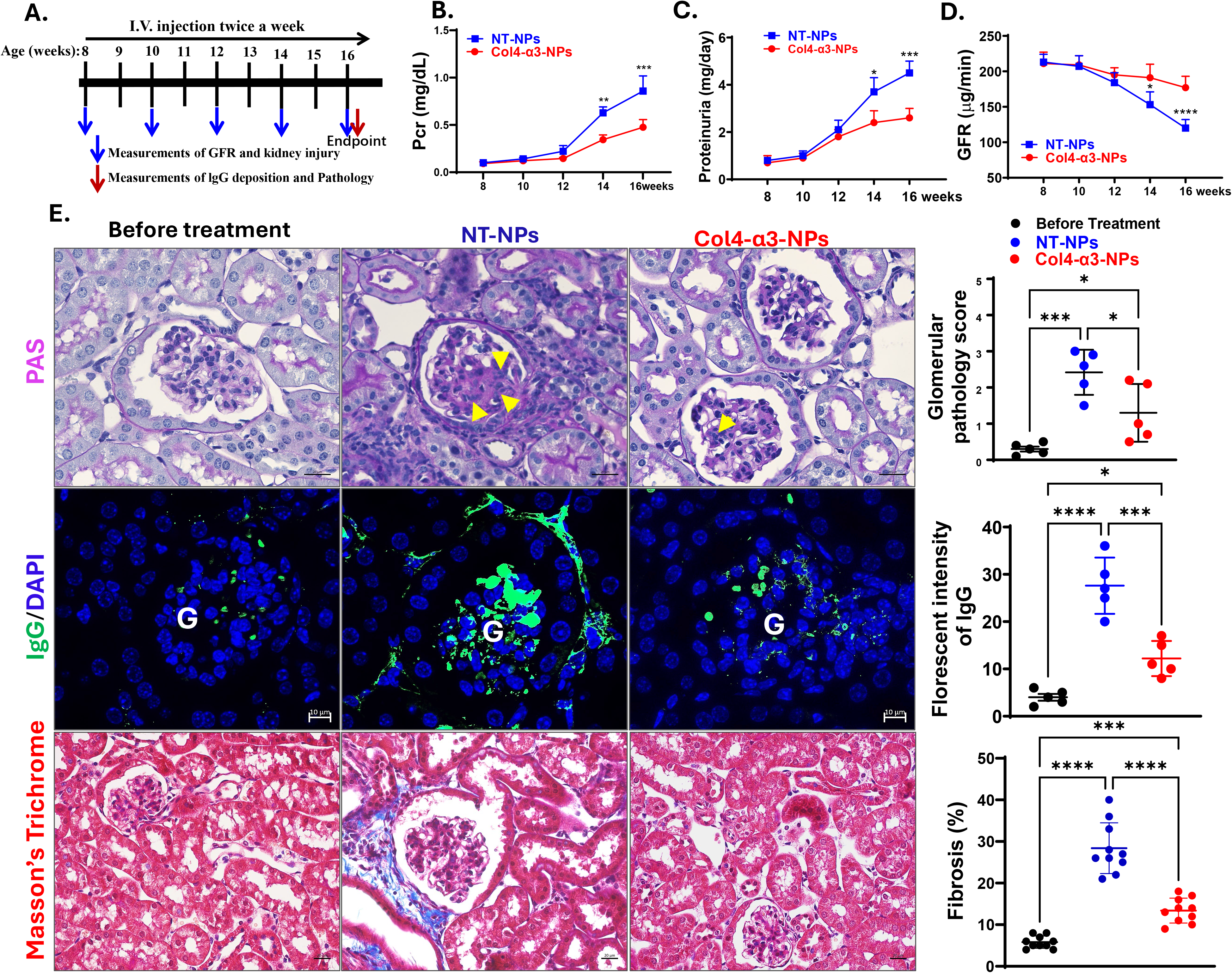
**Targeted delivery of low-dose prednisolone via Col4-**α**3-NPs delayed onset of lupus nephritis.** Lupus-prone MRL-lpr mice were treated with prednisolone encapsulated in either NT-NPs or Col4-α3-NPs, administered twice weekly for 8 weeks. (**A**) Study design. (**B–D**) Plasma creatinine (Pcr), proteinuria, and glomerular filtration rate (GFR) demonstrated significantly improved renal outcomes in the Col4-α3-NP–treated group compared to the NT-NP group, indicating better preservation of kidney function. Statistical analysis was performed using two-way ANOVA followed by Sadak’s multiple comparisons test. N=5 mice/group, *P<0.05; **P<0.01; ***P<0.001, ****p<0.0001 (**E**) Representative kidney sections stained with PAS, Masson’s trichrome, and IgG immunofluorescence show that treatment with prednisolone-loaded Col4-α3-NPs markedly reduced collagen deposition, IgG accumulation, and renal fibrosis. Scale bars represent 20 µm for PAS and Masson’s trichrome staining and 10 µm for IgG immunofluorescence staining. Statistical analysis was performed using One-way ANOVA followed by Tukey’s multiple comparisons test. *P<0.05; ***P<0.001, ****p<0.0001, n=5 mice/group.

After eight weeks, mice treated with prednisolone-loaded Col4-α3-NPs exhibited a 42% lower proteinuria and a 48% lower in Pcr compared to control group (Fig. 5B-C). Additionally, GFR was preserved, showing a 33% higher value in the treatment group (Fig. 5D). These findings suggest that targeted delivery of prednisolone via Col4-α3-NPs effectively ameliorates kidney injury and preserves renal function.

Histopathological analysis revealed that control mice displayed pronounced glomerular hypertrophy and mesangial proliferation, as evidenced by PAS staining (Fig. 5E). In contrast, mice treated with prednisolone-loaded Col4-α3-NPs showed markedly reduced glomerular alterations. Immunofluorescence staining indicated substantial IgG deposition in the glomeruli of NT-NPs group, which was significantly diminished in the Col4-α3-NPs group. Furthermore, Masson’s Trichrome staining demonstrated marked reduction in fibrosis in the kidneys of Col4-α3-NP–treated mice compared with the NT-NP group.

### Potential mechanisms of the Col4-**α**3-NPs loaded with prednisolone for LN

To explore the underlying mechanisms of local treatment versus systemic effects, we isolated immune cells from kidneys, renal lymph nodes and blood at the end of experiments for flow cytometry measurements. As shown in Fig 6, treatment with prednisolone-loaded Col4-α3-NPs in MRL-lpr mice induced changes locally as anticipated. The frequencies of activated T and B cells in the renal lymph nodes (Fig. 6 A–C) were significantly reduced following the Col4-α3-NPs loaded with prednisolone, and the proinflammatory cytokine-producing CD4⁺ T cells (Fig.6 D–F) were markedly decreased in the Col4-α3-NPs treated group than NT-NPs group. Some of the parameters are not statistically significant but show the right trend and will likely reach significance with longer-term treatment (Fig. 6 G). Although, the Col4-α3-NPs treated mice had a higher proportion and number of DCs and macrophages compared to NT-NPs (Fig. 7 A-D), the expression of the co-stimulatory molecules CD80 and CD86 was remarkably attenuated (Fig. 7 E) suggesting a tolerogenic phenotype. Interestingly, the numbers of Tregs were higher in the kidneys, but lower in lymph nodes in Col4-α3-NPs treated mice compared to the NT-NPs controls (Fig. 7 F). These data suggest that either there is a local effect of the treatment in the kidneys resulting in higher Treg enrichment locally in the kidneys due to their migration from lymph nodes to the kidneys or a local induction of Tregs due to the greater tolerogenic phenotype of the DC (61, 62).

**Figure 6.**
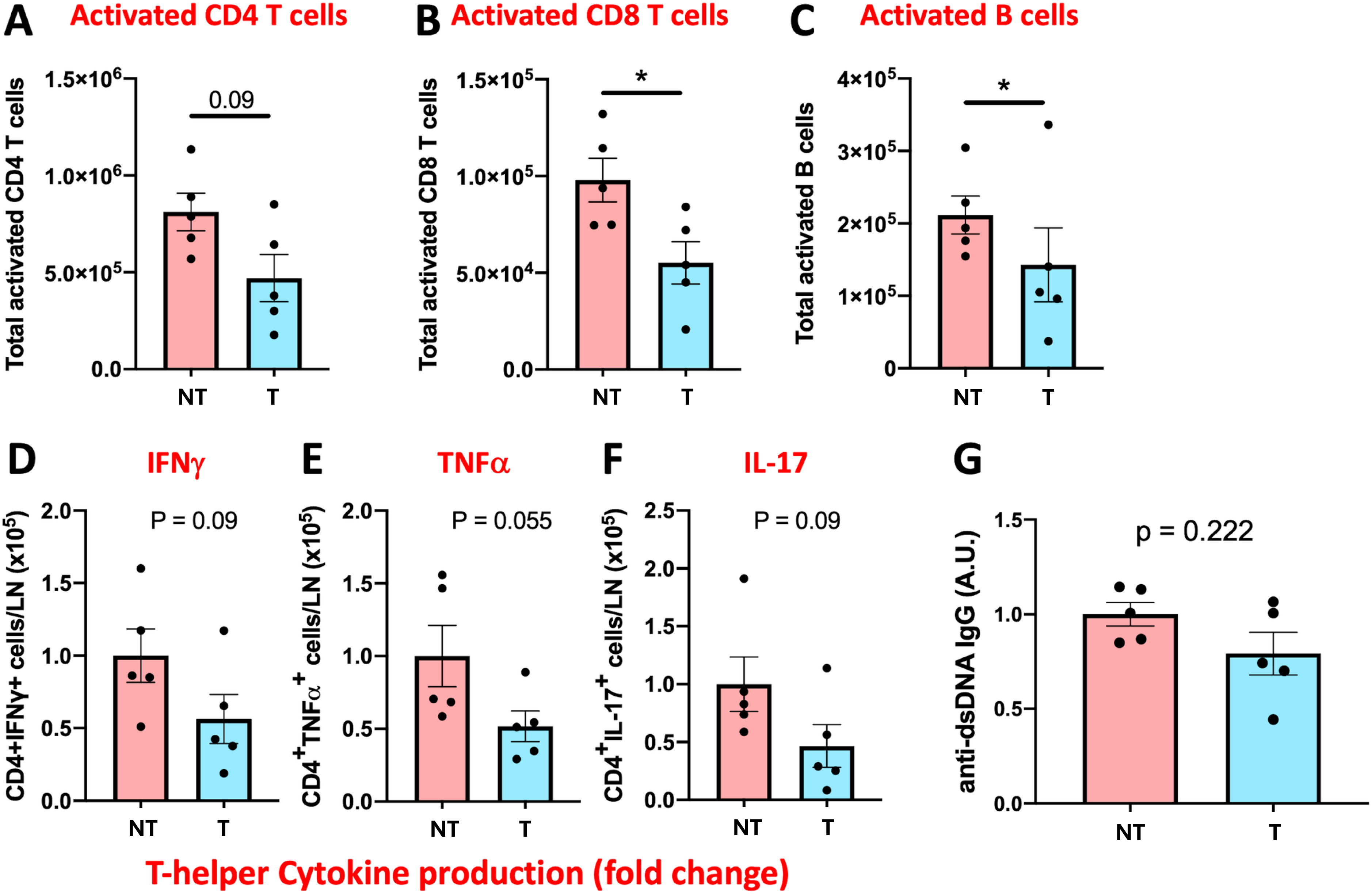
**Immune activation and proinflammatory responses were attenuated in the kidneys of Col4-**α**3-NP–treated MRL-lpr mice.** Treatment with Col4-α3-NPs resulted in reduced frequencies of activated T and B cells in the lymph nodes (**A–C**), and a trend toward decreased proinflammatory cytokine-producing CD4⁺ T cells (**D–F**). However, circulating levels of anti-dsDNA antibodies were only minimally and non-significantly affected (**G**). Statistical analysis was performed using unpaired 2-tailed Student’s t test. *p <0.05, n=5mice/group. NT, NT-NPs; T, Col4-α3-NPs.

**Figure 7.**
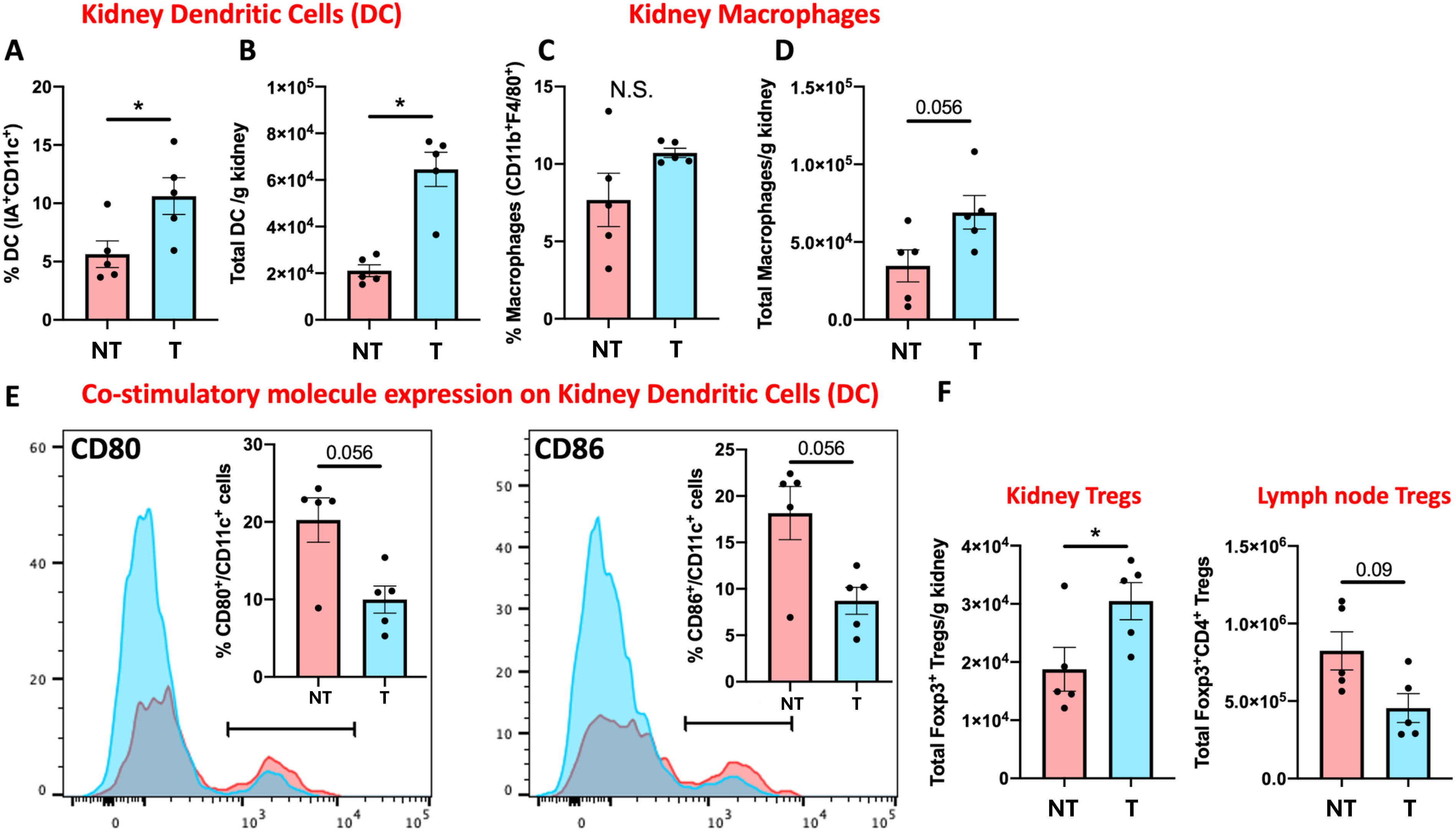
**Kidneys of MRL-lpr mice treated with Col4-**α**3-NPs had a higher proportions and numbers of dendritic cells (DC) and macrophages.** (A-D). The expression of costimulatory molecules CD80 and CD86 on the DC from the Col4-α3-NPs treated mice was lower than the NT-NPs mice. (E) and (F) The number of Tregs was higher in the kidneys of the Col4-α3-NPs-treated mice whereas the number of Tregs in the lymph nodes of Col4-α3-NP-treated mice was lower than the NT-NPs mice. Statistical analysis was performed using unpaired 2-tailed Student’s t test. *p <0.05, n=5mice/group.

## Discussion

In this study, we developed and validated a novel glomerulus-targeted drug delivery system using collagen IV-binding peptide-conjugated liposomal nanoparticles (Col4-α3-NPs) for the treatment of LN. Our findings demonstrate that Col4-α3-NPs exhibit selective glomerular accumulation, extended drug release, and significant therapeutic efficacy with minimal systemic exposure when loaded with prednisolone. This targeted nanomedicine approach offers a promising strategy for preventing or treating glomerular diseases such as LN, where site-specific delivery can improve therapeutic outcomes while reducing off-target systemic toxicity.

The development of Col4-α3-NPs represents a significant step toward precision therapy for glomerular diseases. The pathogenesis of LN involves immune complex deposition and inflammatory injury in the glomeruli, making them a logical target for therapeutic intervention (3, 63–65). However, existing systemic treatments, particularly glucocorticoids (GCs), are associated with broad immunosuppression and substantial adverse effects (7, 66, 67). By leveraging the unique exposure of GBM collagen IV to the circulation through fenestrated endothelium, we engineered a delivery system capable of precise glomerular targeting. Our *in-silico* modeling and SPR analysis confirmed strong and specific binding affinity of the selected peptide to Col4-α3, a major GBM component, with minimal cross-reactivity to Col4-α1.

The biodistribution studies in mice confirmed that Col4-α3-NPs preferentially accumulate in glomeruli with limited off-target deposition in other organs. Liposomal NPs that are administered intravenously, enter the bloodstream quickly and spread throughout the body. The initial distribution phase is marked by the accumulation of Liposomal NPs to various tissues. However, a significant amount tends to accumulate in the liver and spleen due to the filtering and phagocytic functions of these organs(68, 69). The reticuloendothelial system, including macrophages, is vital for capturing these liposomes. Depending on their formulations such as size, charge, and surface modifications—some Liposomal NPs can minimize the detection and clearance by the immune system, allowing their longer circulation in the bloodstream. For example, the pegylated Liposomal NPs used herein are known to have extended circulation times, which improves their distribution to other tissues, including the kidneys. Over time, Liposomal NPs are taken up by the kidneys due to the presence of targeting agents. We did not detect strong signals in the heart or lungs, which is consistent with the well-known low accumulation of Liposomal NPs in these organs. This degree of tissue specificity surpasses previous NP strategies that often relied on passive targeting via particle size or charge (28, 29, 70, 71). Importantly, the targeted delivery did not compromise renal hemodynamics or induce structural injury, confirming the biocompatibility and safety of the Col4-α3-NPs. A key advantage of our system lies in its prolonged and controlled release properties. Both *in vitro* and *in vivo* experiments demonstrated that the encapsulated drug—whether modeled by calcein or FITC-steroid—was released over 24 to 48 hours in a stable and extended manner, supporting the feasibility of reduced dosing frequency. This extended-release profile likely contributes to improved drug efficacy and reduced systemic exposure, which are particularly beneficial in chronic conditions like LN.

Another major goal of targeting specifically the glomeruli is that it enables the induction of a tolerogenic immune environment in the kidney, potentially reducing the need for aggressive systemic immunosuppression. Recent mechanistic studies have identified the role of tertiary lymphoid structures (TLS) in sustaining local autoantibody production and antigen presentation, making them a target for disrupting renal-specific immune activation (72). Also, tissue-resident memory T cells (TRM), particularly CD8+ subsets, contribute to persistent inflammation and are modulated through pathways such as NLRP3 inflammasome inhibition. Cytokine signaling, notably IL-18 and IFN-γ, drives Th1-dominant responses in LN kidneys, and local blockade of these pathways has shown promise in reducing renal pathology. Together, these insights underscore the feasibility and importance of local immune modulation in LN, paving the way for therapies that preserve systemic immune integrity while effectively controlling renal inflammation.

To this end, we have shown that in lupus-prone MRL-lpr mice, treatment with prednisolone-loaded Col4-α3-NPs significantly improved renal outcomes compared to non-targeted controls. Proteinuria, plasma creatinine, and GFR were markedly improved, and histological examination revealed reduced glomerular hypertrophy, IgG deposition, and fibrosis. Notably, therapeutic efficacy was achieved with a substantially lower prednisolone dose (0.34 mg/kg) than the conventional systemic dose used to treat lupus (5–10 mg/kg)(57–59, 73), highlighting the advantage of glomerular-targeted drug delivery. Based on the well-established therapeutic dose range and our preliminary findings (Figure S3), free prednisolone at the same low dose was not expected to provide meaningful therapeutic benefit and was therefore not included in the main efficacy study. The primary objective of this study was to determine whether glomerular-targeted nanoparticle delivery could achieve therapeutic efficacy with markedly reduced glucocorticoid exposure.

Further, our findings suggest that the Col4-α3-NPs exert both anti-inflammatory and immunomodulatory effects at the site of injury. Flow cytometry revealed an increase in renal regulatory T cell (Treg) populations and a reduction in costimulatory molecule expression (CD80/CD86) on local antigen-presenting cells, indicative of a more tolerogenic immune environment. Since CD80 and CD86 are well-established markers of antigen-presenting cell activation, their reduced expression suggests that these immune cells are in a less activated, more tolerogenic state rather than a pro-inflammatory phenotype(74). This correlates with the reduced proportion of activated T cells and lower production of proinflammatory cytokines IFNγ, TNFα and IL-17 by T-cells. Approaches to enhance Tregs have been shown to be promising to provide durable remission in both experimental (75) and clinical studies (76) in SLE and lupus nephritis (77). The concurrent reduction of Tregs in lymph nodes suggests a redistribution or localized induction of Tregs in the kidneys as a mechanism underlying efficacy of our targeted therapy. Local prednisolone hampered the LPS induced upregulation of co-stimulatory molecules and limited the expression of IL-12 and IL-23 along with improved Treg function (45, 75, 78, 79). Localized stimulation with prednisone upregulated FOXP3 and STAT5 in tissue resident decidual immune cells (80) as well as induced robust Tregs expansion in vitro (81). On the other hand, systemic high-dose methylprednisolone in SLE patients only showed a transient increase in Tregs that rapidly decline (82). In the present study, Col4-α3-NP treatment produced a modest, but not statistically significant, reduction in circulating anti-dsDNA antibody levels. This may reflect the long half-life of existing autoantibodies and persistence of antibody-producing plasma cells, which could mask treatment-related changes in newly generated autoantibodies(83). Nevertheless, Col4-α3-NP significantly reduced the proportion of activated B cells (Fig. 6). The limited effect on circulating anti-dsDNA antibodies is also consistent with the proposed mechanism of Col4-α3-NP, which preferentially delivers prednisolone to the kidney and primarily suppresses local immune complex-mediated renal inflammation rather than systemic autoantibody production. Accordingly, treatment significantly reduced immune complex deposition and improved renal pathology and kidney function despite only modest changes in circulating anti-dsDNA levels. It is also possible that treatment altered the pathogenicity rather than the total amount of anti-dsDNA antibodies. In a previous lupus nephritis study, increased Treg activity protected the kidney without significantly reducing total anti-dsDNA antibody levels, but significantly decreased the ratio of pathogenic IgG2a to non-pathogenic IgG2b anti-dsDNA antibodies(75, 84). Thus, a localized immunoregulation could contribute to disease attenuation without the systemic immunosuppression seen with free GCs—an effect that warrants further investigation.

Compared to previous kidney-targeting strategies—such as those employing antibodies, aptamers, or charge-based accumulation—our Col4-α3-NPs system offers superior specificity and favorable pharmacokinetics. Additionally, the use of clinically relevant materials (liposomes and prednisolone) enhances the translational potential of this platform. However, there are limitations to this study. While the current results are compelling, they are limited to murine models and short-term outcomes. Additionally, the treatment effects of prednisolone-loaded Col4-α3-NPs on well-established lupus nephritis were not evaluated, as the Col4-α3-NPs was administered at the initiation of the development of lupus nephritis in mice, however, MRL/lpr mice do accumulate circulating autoantibodies, indicative of an SLE like disease soon after weaning (85) and immune complex deposition can be detected by the time we initiated the nanoparticle treatment(60). Evaluation of therapeutic efficacy in mice with established lupus nephritis will be the focus of future studies and represents an important step toward clinical translation. Additionally, the glomerular localization of Col4-α3-NPs was not directly evaluated in MRL/lpr mice. Although glomerular basement membrane remodeling in lupus nephritis may increase the accessibility of the Col4-α3 epitope, this remains to be experimentally confirmed. Future studies will characterize the intrarenal biodistribution and glomerular localization of Col4-α3-NPs in established lupus nephritis. In addition, further investigation is needed to optimize dosing, evaluate long-term safety, and determine potential immune responses to the peptide-liposome formulation. Finally, more comprehensive immune profiling, including macrophage polarization, dendritic cell subsets, regulatory T cells, and anti-inflammatory cytokines, will help define the immunomodulatory mechanisms of Col4-α3-NPs. Validation of therapeutic efficacy in additional lupus nephritis classes and in male mice will also be important for assessing the broader translational potential of this targeted therapeutic strategy. Furthermore, evaluating the platform’s versatility with other therapeutic agents or for co-delivery strategies (e.g., combination therapy with immunomodulators or biologics) could expand its clinical utility.

In conclusion, our study demonstrates a precision Col4-α3-NP-targeted therapeutic strategy, addressing a major unmet need in LN. By combining glomerular specificity, prolonged drug release, and immunomodulatory potential, this platform addresses the critical need for site-specific, efficacious, and safer treatment options in LN and potentially other glomerular diseases.

### Clinical perspectives

Background: LN is a serious complication of Systemic Lupus Erythematosus that often requires long-term treatment with systemic immunosuppressive drugs, especially glucocorticoids, which can cause significant side effects. Although nanoparticle-based drug delivery systems have advanced in recent years, effective glomerulus-specific targeting remains challenging. In this study, we developed a collagen IV-α3-targeted liposomal nanoparticle system designed to selectively deliver prednisolone to the glomeruli.

Results: We demonstrated that Col4-α3-NPs preferentially accumulated in kidney glomeruli and provided sustained drug release. In lupus-prone mice, prednisolone-loaded Col4-α3-NPs significantly improved kidney function, reduced proteinuria, IgG deposition, and renal fibrosis, and increased renal regulatory T cell populations. Importantly, these beneficial effects were achieved using a substantially lower prednisolone dose compared with conventional systemic therapy.

Significance: Our findings suggest that glomerulus-targeted nanoparticle therapy may provide a more effective and safer treatment strategy for lupus nephritis by reducing systemic steroid exposure while preserving therapeutic efficacy. This targeted delivery strategy may also be adaptable for the treatment of other glomerular diseases.

## Availability of data and materials

All the data and materials supporting the findings of this study are available within the article. Further enquiries can be directed to the corresponding author.

## Author contributions

LW, EM, SSM, SM and RL design the project and draft the manuscript. LW, EM, RV, RS, CE, AP, WL, AM and KM performed experiments, analyzed data. EM, AP, WL, AM and KM designed, synthesized and characterized the nanoparticles. LW, CE, KW, JW, and JZ performed cell culture, animal experiments, histology and data analysis. LF performed histological analysis. LW, EM, SM, SSM, and RL edited the manuscript. SM and RL supervised the study.

