## Supplemental for "Glomerulus-Targeted Nanotherapy via Collagen IV-α3 Binding Enhances Renal Immunoregulation in Lupus Nephritis"

**The file includes:**

**Figs. S1, S2, S3 and Table S1.**

**Figure S1.**

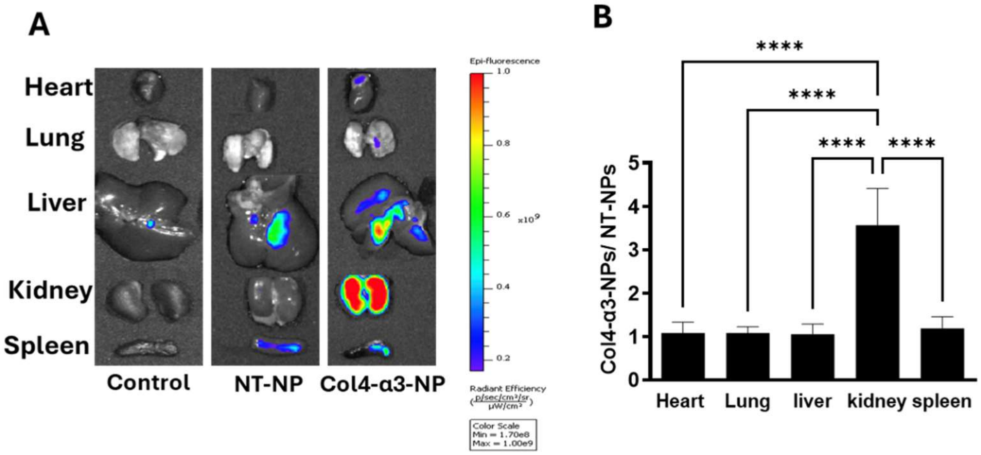


**Figure S1. Nanoparticles showed Col4-α3-NPs preferentially accumulated in kidneys. A** Representative image of mouse organs collected 48 hours after injection with Saline, rhodamine-labeled NT-NPs or Col4-α3-NPs. **B.** Quantification of fluorescence signal intensity, expressed as the ratio of Col4-α3-NPs to NT-NPs. Statistical analysis was performed using unpaired 2-tailed Student's t test, n=5 mice/group, ****$p<0.0001$.

**Figure S2.**

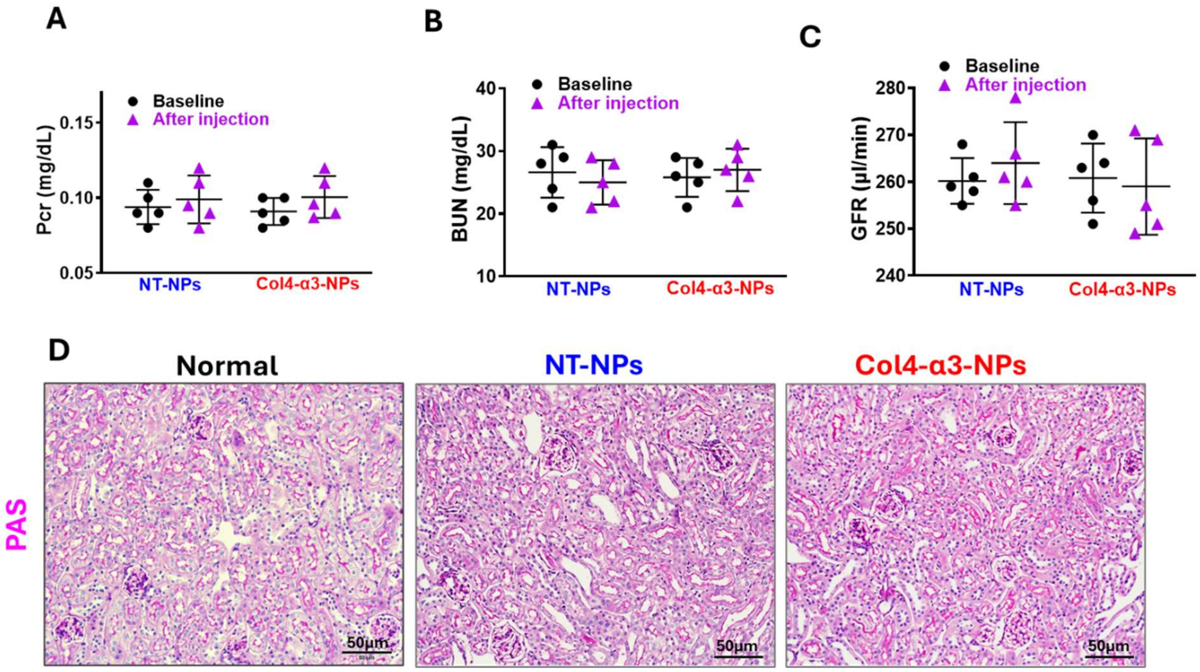


**Figure S2.** Nanoparticles exhibit no observable side effects in Mice. NT-NPs and Col4-α3-NPs were administered intravenously to C57BL/6 mice for 8 weeks. Plasma creatinine (Pcr), blood urea nitrogen (BUN), glomerular filtration rate (GFR), and renal histology were assessed before and 24 hours after nanoparticle (NP) administration. No significant changes in Pcr (A), BUN (B), or GFR (C) were observed post-injection. Renal blood flow (RBF), measured following GFR assessment, remained unchanged. (D)Periodic acid–Schiff (PAS) staining revealed no morphological alterations or signs of kidney injury. Statistical analysis was performed using unpaired 2-tailed Student's t test, n=5 mice/group.

**Table S1. Biochemical Parameters**

| Parameters | Col4-α3-NPs | NT-NPs | Significance |
|---|---|---|---|
| HCO3- act (mmol/L) | 18.5±1.3 | 19.2±1.4 | ns |
| Na+ (mmol/L) | 139.6±0.4 | 140.5±0.5 | ns |
| K+ (mmol/L) | 4.15±0.3 | 4.07±0.1 | ns |
| Ca++ (mmol/L) | 1.28±0.2 | 1.29±0.1 | ns |
| Cl- (mmol/L) | 107.8±0.4 | 108.7±0.3 | ns |
| Glucose (mg/dL) | 220.2±33.4 | 216.67±15.2 | ns |
| Body Weight (g) | 27.6±2.1 | 28.9±1.8 | ns |
| Kidney Weight(g) | 0.146±0.006 | 0.148±0.005 | ns |
| RBF (ml/min) | 0.85±0.07 | 0.79±0.05 | ns |

$HCO_3^-$ act, actual bicarbonate; RBF, renal blood flow; ns, not significant

**Table S1**. presents the key electrolyte parameters, confirming that both the NT-NP and Col4-α3-NP groups maintained electrolyte levels within the normal range. Statistical analysis was performed using unpaired 2-tailed Student’s t test, ns= not significant, n=5 mice/group.

**Figure S3.**

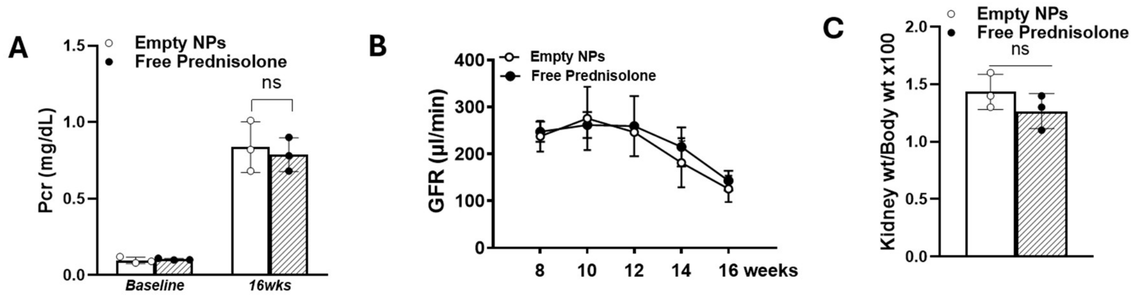


**Figure S3.** Low-dose free prednisolone did not attenuate lupus nephritis progression. In a preliminary study, lupus mice were treated with the same low dose of free prednisolone used in the Col4-α3-NP formulation or with empty nanoparticles (NPs) by intravenous injection for 8 weeks. Plasma creatinine (Pcr) and glomerular filtration rate (GFR) were measured before and at the end of treatment, and the kidney-to-body weight ratio was determined at the study endpoint. No significant differences in plasma creatinine (A), GFR (B), or kidney-to-body weight ratio (C) were observed between the groups. Data are presented as mean ± SEM; n = 3 mice per group.
